# Complete genome of a Measles virus from Roraima state, Brazil

**DOI:** 10.1101/473462

**Authors:** Cátia Alexandra Ribeiro Meneses, Valdinete Alves do Nascimento, Victor Costa de Souza, Rodrigo Melo Maito, Marconi Aragão Gomes, Claudeth Rocha Santa Brígida Cunha, Ilma de Aguiar Antony, Maria Eliane Oliveira e Silva, Daniela Palha de Souza Campos, André de Lima Guerra Corado, Karina Pinheiro Pessoa, Dana Cristina da Silva Monteiro, Osnei Okumoto, Marília Coelho Cunha, Flávia Caselli Pacheco, Felipe Gomes Naveca

## Abstract

Measles is a human infectious disease of global concern caused by the measles virus. In this study, we report the complete genome sequencing of one measles isolate, genotype D8, obtained in Boa Vista city, the capital of the Roraima State, Brazil, directly from the urine sample. Phylogenetics reconstruction grouped the genome described in this study with samples from Australia, Italy, United Kingdom, and the USA. To our knowledge, this is the first complete genome of a wild-type measles virus from Latin America. Therefore, the present data strengthens the current knowledge about the molecular epidemiology of measles worldwide.

**Sponsorships:** CNPq / CAPES / MS-DECIT / Fiocruz

---

Measles is a highly contagious airborne disease that begins as an acute febrile illness with fever, coryza, conjunctival hyperemia and cough, which are accompanied by a maculopapular skin rash. Some measles patients, mainly those immunocompromised, may evolve to severe complications including blindness and life-threating forms as severe diarrhea and pneumonia. Two highly severe, but rare neurologic forms, are also related to measles virus infection, the measles inclusion-body encephalitis MIBE and the subacute sclerosing panencephalitis (SSPE) (1).

The measles virus is an enveloped, single-stranded, negative-sense RNA virus, with a genome around 16Kb that belongs to the *Paramyxoviridae* family, genus *Morbillivirus*. A total of eight proteins, six structural (nucleoprotein, phosphoprotein, matrix, fusion, haemagglutinin, and the large polymerase protein) and two non-structural (V and C proteins) are encoded by the viral genome (1). Due to the lack of proofreading activity of RNA polymerases and the higher viral titers achieved during the acute phase of illness, RNA viruses are more likely to accumulate variability in their genome’s sequences, in comparison to DNA viruses (2). Regarding the measles virus, this diversity is reflected in 24 genotypes known up to date (A, B1, B2, B3, C1, C2, D1, D2, D3, D4, D5, D6, D7, D8, D9, D10, D11, E, F, G1, G2, G3, H1, and H2), but from October 2017 to September 2018, only five genotypes (B3, D4, D8, D9, and H1) were reported (http://www.who-measles.org/Public/Data_Mnt/who_map.php).

Measles is a vaccine-preventable disease which has decreased mortality over the last decades. Since the 1980s, the total number of deaths related to measles illness dropped from more than 2 million to approximately 100,000 per year due to the improvement of social indicators (e.g., better nutrition), as well as the establishment of the global efforts to increase vaccination coverage (1). Nevertheless, timely surveillance of suspected measles cases with highly sensitive and specific molecular diagnostic tools, together with genetic characterization of isolates, is of paramount importance for the global efforts of virus elimination (1, 3).

In August 2018, the Central Laboratory of the Roraima State (LACEN-RR) started testing measles suspected samples using the Real-Time PCR protocol developed by CDC-USA. Procedures followed the annex 6.2 of the Manual for the Laboratory-based Surveillance of Measles, Rubella, and Congenital Rubella Syndrome (Third edition, June 2018) freely available at http://www.who.int/immunization/monitoring_surveillance/burden/laboratory/Annex_6.2.pdf?ua=1. Since then, 172 samples were tested, and 39 were positive.

Thus, as a request of both Roraima’s state and the Brazilian health surveillance authorities, we select one positive sample with high viral load (Ct 22.0) to submit for a protocol for the entire genome amplification and nucleotide sequencing. Initially, we aligned all measles D8 genomes with MAFFT v7.388 (4). The consensus sequence was used to design primers over the entire genome of measles D8 with the aid of Primal software (5) spanning around 400bp distances. Two other primers targeting the initial 28 bases of the 5’ UTR region and the final 25 bases of the 3’ UTR were designed with a modified version of Primer3 v2.3.7 embedded in Geneious software v10.2.6 (6). This primer design may be used for NGS sequencing, as previously described for Zika virus (5), or capillary sequencing as we performed in this study. Primer sequences and the details for the RT-PCR scheme used for entire genome amplification and nucleotide sequencing are in the supplemental files 1 and 2 of this manuscript.

Briefly, four overlapping amplicons encompassing the entire measles genome were generated using Superscript IV One-Step RT-PCR System (ThermoFisher Scientific). Amplicons were precipitated with molecular biology grade PEG8000 and used as a template for nucleotide sequencing with BigDye terminator v3.1 in an ABI 3130 genetic analyzer.

A total of 63 trace files were trimmed for quality and used for assembly employing Geneious's map to reference tool and the NCBI measles reference sequence, available in GenBank under accession number NC_001498.1. The final whole genome sequence of MVs/Roraima.BRA/31.18[D8] isolate contains 15,894 nucleotides with no ambiguity, or unidentified (N) bases, a Q40 score of 99.8% and a GC content of 47.4 %.

Initially, the sequence reported here was genotyped using both the Nucleoprotein (N) gene and the Hemagglutinin (H) gene with the measles genotyping tool available at http://www.who-measles.org/Public/Tool/genotype_tool.php, and the D8 genotype was confirmed. Subsequently, we conducted a maximum likelihood phylogenetic analysis with PhyML (7) using the N gene region of the Roraima sample, the genotypes reference sequences available at MeaNS http://www.who-measles.org/ and four other sequences representing the genotype D8 lineages (FJ765078.1 – MVi/Villupuram.Ind/03.07; JX486001.1 - MVi/Hulu-Langat.MYS/26.11; KF683445.1 - MVs/Frankfurt_Main.DEU/17.11; KT588030.1 - MVs/Republic_of_Komi.RUS/35.13). This approach confirmed that the Roraima sample belongs to the lineage MVi/Hulu-Langat.MYS/26.11 (Supplemental file 3).

Secondly, a BLAST search was conducted using the megablast algorithm against the entire NR database. The first five closest matches are samples from Brisbane, Australia (MH638233, score 29137, 15834/15862 identical sites); Ancona, Italy (MH173047, score 29000, 15816/15872 identical sites); California, USA (KY969480, score 28755, 15773/15874 identical sites); Virginia, USA (JN635404, score 28731, 15782/15894 identical sites) and London, England (KT732231, score 28476, 15592/15678 identical sites).

Finally, a dataset containing all 42 complete measles genomes belonging to the genotype D8 available in GenBank in 06-Nov-2018, and the sequence obtained in this study, were analyzed with jModeltest 2.1.7 v20150220 and likelihood scores were computed for five substitutions schemes (40 models). The GTR+I model was selected by AICc and used for Bayesian phylogenetic reconstruction using two runs with 20mi generations with MrBayes 3.2.6 (8). This analysis showed two main clades, one containing 31 sequences from the United Kingdom 2012-2013, one from Netherlands (2013), one from Canada (2010), one from Germany (2013), two from Vietnam (2014), and one sequence from Texas, USA (2007), as the outgroup. The second clade contains the same five sequences returned from the BLAST analysis with the highest scores, from Italy (2017), Australia (2015), United Kingdom (2012), and two from USA, California (2010) and Virginia (2009), been this last one the outgroup of this clade. Furthermore, the Bayesian tree showed that the isolate MVs/Roraima.BRA/31.18[D8] grouped in this second clade, close together with the sample from Brisbane, Australia, collected in 2015 and the sample from Ancona, Italy collected in November 2017, with a high posterior probability (1.0) (Figure).

**Figure.**
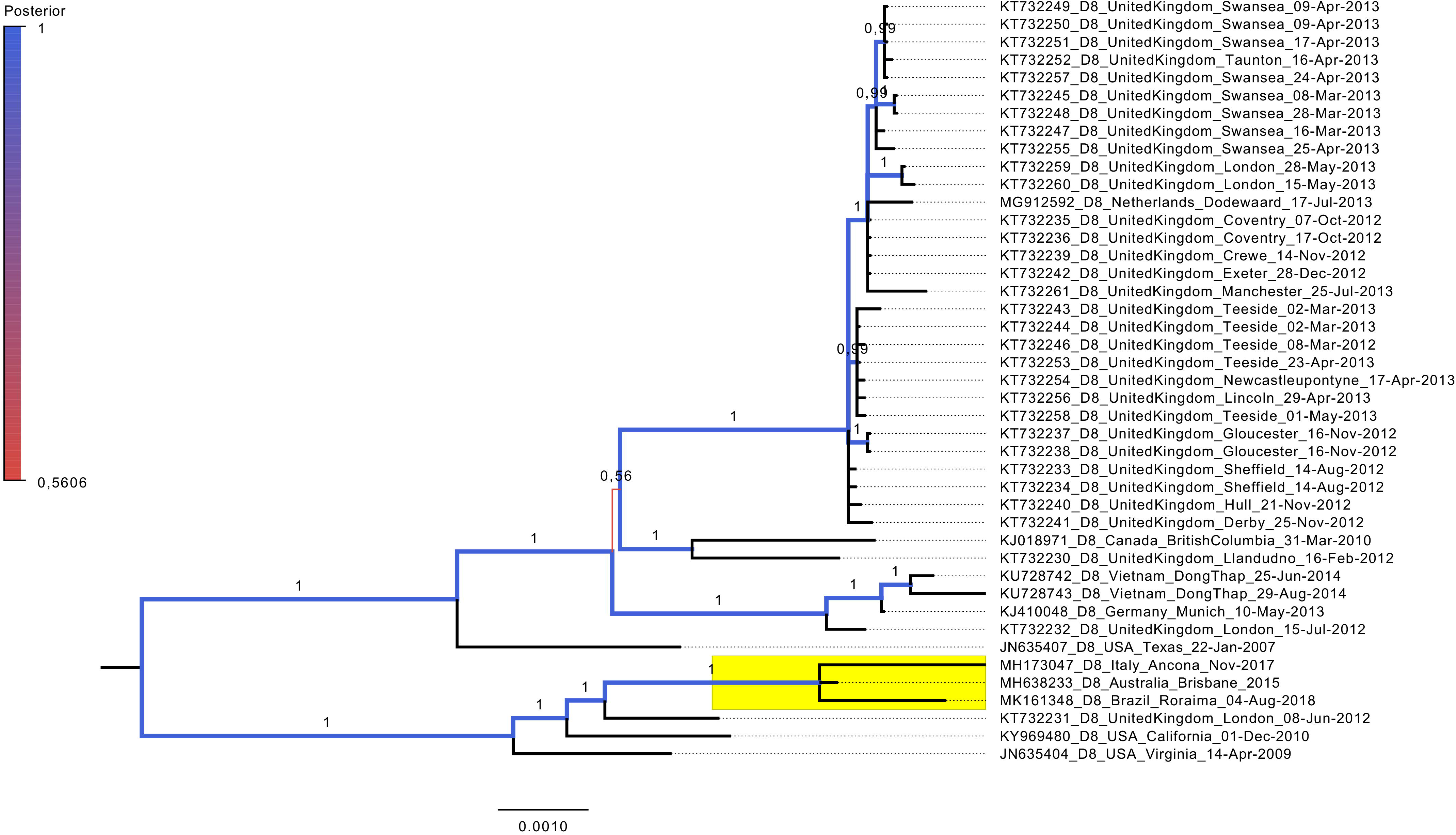
Phylogenetic tree of complete measles virus genomes. A mid-rooted Bayesian tree with increasing node order was constructed with MrBayes 3.2.6 and 43 taxa (complete CDS and intergenic regions from position 108 to 15,785 related to the reference sequence NC_001498.1) representing all the complete measles (genotype D8) genomes available at GenBank in 06-Nov-2018 and the sequence reported in this study. Branches are colored by posterior probabilities, according to the legend, and the specific posterior probabilities values are shown. The clade containing the sequence described in this study is highlighted in yellow. The scale bar represents nucleotide substitutions per site.

Surprisingly, we did not find any measles complete genome record from Latin America on the public databases including GenBank, Virus Pathogen Resource (ViPR - http://www.viprbrc.org), and MeaNS. This fact hindered a more detailed analysis of the complete genome reported in this study in the context of the local transmission, strengthening the necessity of sequencing more fully genomes from this region.

Previous studies in Brazil have primarily concentrated on genotyping a small fragment of 450bp of the measles nucleoprotein (9,10). Although, these studies have undoubtedly contributed to the molecular epidemiology of measles, sequencing larger genome fragments or better yet, complete viral genomes, is pivotal for a better understanding of the epidemic dynamics of any emerging or remerging viral diseases (11,12).

## Nucleotide sequence accession number

The complete genome sequence of the MVs/Roraima.BRA/31.18[D8] isolate is available at GenBank under the accession number MK161348.

## ACKNOWLEDGMENTS

FGN is funded by Fundação de Amparo à Pesquisa do Estado do Amazonas – FAPEAM (http://www.fapeam.am.gov.br, call 001/2013 – PPSUS / 062.00656/2014 and call 001/2014 – PROEP / 062.01939/2014); Conselho Nacional de Desenvolvimento Científico e Tecnológico (http://www.cnpq.br, grant 440856/2016-7) and Coordenação de Aperfeiçoamento de Pessoal de Nível Superior (http://www.capes.gov.br, grants 88881.130825/2016-01 and 88887.130823/2016-00) call MCTIC/FNDCT-CNPq / MEC-CAPES/ MS-Decit 14/2016 – Prevenção e Combate ao vírus Zika. The authors thank the Program for Technological Development in Tools for Health-PDTIS FIOCRUZ for use of nucleotide sequencing facilities at ILMD—Fiocruz Amazônia. The funders had no role in study design, data collection and analysis, decision to publish, or preparation of the manuscript.

---

**Supplemental file 1**. Primers designed to amplify and to sequence the entire genome of measles Genotype D8.

**Supplemental file 2**. Primers sets used to amplify and to sequence the entire measles genome. Four amplicons were generated: Amplicon 1 (primers 5UTR + 11R, 3614bp); Amplicon 2 (11L + 26R, 5003bp); Amplicon 3 (24L + 40R, 5405bp); Amplicon 4 (37L + 3UTR, 4682bp). Each amplicon was sequenced with the primers listed below each scheme.

**Supplemental file 3**. Phylogenetic tree of measles genotypes based in the N gene. A mid-rooted ML tree with increasing node order was constructed with PhyML online server available at http://www.atgc-montpellier.fr/phyml/. All 24 genotypes and the four lineages of genotype D8 are represented. The D8 sequences are grouped in the blue clade, whereas the clade containing the Roraima sample is highlighted in yellow.

